# The Long Pentraxin 3 (PTX3) Suppresses Immunity to Cutaneous Leishmaniasis by Negatively Regulating Th17 Response

**DOI:** 10.1101/585315

**Authors:** Gaurav Gupta, Ping Jia, Rohit Sharma, Romaniya Zayats, Sayonara M. Viana, Lianyu Shan, Zhriong Mou, Aldina Barral, Viviane S. Boaventura, Thomas T. Murooka, Abdel Soussi-Gounni, Camila I. de Oliveira, Jude E. Uzonna

## Abstract

The long Pentraxin 3 (PTX3), a soluble pattern recognition molecule, plays a critical role in inflammation, tissue repair and wound healing. Here, we show that PTX3 regulates disease pathogenesis in cutaneous leishmaniasis (CL). PTX3 expression is increased in active skin lesions in patients and mice during CL, with higher levels being expressed in individuals with severe disease. PTX3 deficient (PTX3^-/-^) mice were highly resistant to *L. major* infection and the enhanced resistance was associated with increased IL-17 response. Neutralization of IL-17A abolished this enhanced resistance while treatment with recombinant PTX3 resulted in reduced IL-17A response and increased susceptibility to *L. major* infection. Naïve CD4^+^ T cells from PTX3^-/-^ mice displayed increased differentiation into Th17 cells, which was reversed in the presence of recombinant PTX3. The enhanced Th17 response observed in PTX3^-/-^ cells was associated with increased *Leishmania* specific IL-6 production from dendritic cells along with enhanced expression of Th17-specific transcription factors including RORγt, AhR and STAT3. Addition of recombinant PTX3 significantly inhibited the expression of Th17-specific transcription factors and dramatically reduced the frequency of Th17 cells in Th17-polarizing cultures of PTX3^-/-^ CD4^+^ T cells. Collectively, our results show that PTX3 contributes to the pathogenesis of CL by suppressing Th17 differentiation and IL-17A production.

**Author Summary:** Cutaneous leishmaniasis (CL) is caused by several species of *Leishmania*. Currently, there is no approved vaccine against human CL because of the poor understanding of the mechanisms that regulate disease pathogenesis and correlates of protective immunity. Because the long pentraxin 3 (PTX3, a soluble pattern recognition molecule that forms an integral part of the host innate immunity), regulates inflammation and tissue repair, which are critical physiological events associated with resolution of skin lesions during CL, we investigated its role in disease pathogenesis.

Here, we show that PTX3 levels were elevated in skin-lesions in patients and mice during CL. Using a loss of function approach, we showed that PTX3 contributes to pathogenesis, and this was associated with increased IL-17A responses. Neutralization and recombinant cytokine treatment studies showed that the increased resistance of PTX3 deficient mice to *L. major* is due to enhanced Th17 response in these mice. We further show that PTX3 negatively regulates IL-6 production by dendritic cells and the expression of IL-17A-specific transcription factors (including RORγT, STAT3, IRF4, BATF and AhR) in CD4^+^ T cells. Collectively, these findings show that PTX3 is a negative regulator of Th17 response and protective immunity during *L. major* infection.

## Introduction

Cutaneous leishmaniasis (CL) is caused by several species of protozoan parasites that belong to the genus *Leishmania.* The disease is endemic to Middle East, Asia, Latin and Central America and North Africa (*1*). Resistance to CL is usually associated with the development of strong IFN-γ–producing CD4^+^ Th1 cells, which activate macrophages to produce nitric oxide (NO), an effector molecule for killing intracellular parasites (2-5). In contrast, susceptibility has been associated with IL-4 and IL-10 production by Th2 cells, which are cytokines that deactivate macrophages and inhibit their ability to kill intracellular parasites (2, 6). Besides Th1 and Th2 cells, IL-17A-secreting Th17 cells have also been shown to mediate either host protection (7-10) or susceptibility (11-13) to leishmaniasis. However, the series of events that leads to the induction of Th17 responses in CL are unknown. Our group recently showed that Th17 activation and IL-17A production during allergic asthma was regulated in part by the long Pentraxin 3 (PTX3) (14), a soluble pattern recognition molecule that forms an integral part of the host innate immunity (15, 16). Whether PTX3 also played a central role in regulating Th17 responses during CL is unknown.

PTX3 is expressed by both immune and non-immune cells, such as myeloid dendritic cells (17, 18), neutrophils (19), macrophages (20), mononuclear phagocytes, endothelial cells (21), smooth muscle cells (22), epithelial cells (23, 24), fibroblasts (25), and adipocytes (26). PTX3 is involved in pathogen recognition (15, 27, 28) and plays an important protective role in bacterial (29, 30), fungal (15, 31) and viral (32, 33) infections by modulating the host inflammatory response. Interestingly, PTX3 is capable of promoting (34) as well as suppressing (35, 36) tissue damage due to excessive inflammation. In addition, PTX3 also participates in wound healing and tissue repair (37).

Given the well characterized roles of PTX3 in inflammation and tissue repair, which are critical physiological events associated with resolution of skin lesions during CL, we investigated whether PTX played a protective or pathogenic role during CL. We show that PTX3 negatively regulates host immunity to *L. major* infection by suppressing Th17 differentiation and IL-17A production by CD4^+^ T cells in infected mice.

## Results

### PTX3 negatively regulates the pathogenesis of cutaneous leishmaniasis

PTX3 has been shown to regulate immunity against a wide range of pathogens (15, 30, 31) and to participate in wound healing and tissue repair (37) by modulating the host inflammatory responses. To determine whether PTX3 contributes to disease pathogenesis in CL, a disease characterized by cutaneous inflammation, we first determined changes in the levels of PTX3 expression during active infections in humans and mice. RT-PCR analysis showed that the expression of PTX3 mRNA was significantly (15-fold, p < 0.01) higher in skin lesion biopsies from *L. braziliensis*-infected patients compared to healthy controls (Fig 1A) and correlates with disease severity such that the levels were highest in individual with disseminated cutaneous leishmaniasis (DCL), the most severe form of the disease (Supplementary Fig 1A). Likewise, the expression of PTX3 mRNA (Fig 1B) and protein (Figs 1C and D) was higher at the site of *L. major* (which also causes CL akin to *L. braziliensis*), infection in mice compared to uninfected sites and the expression was mostly restricted to CD68^+^ cells (Fig 1D).

**Fig 1:**
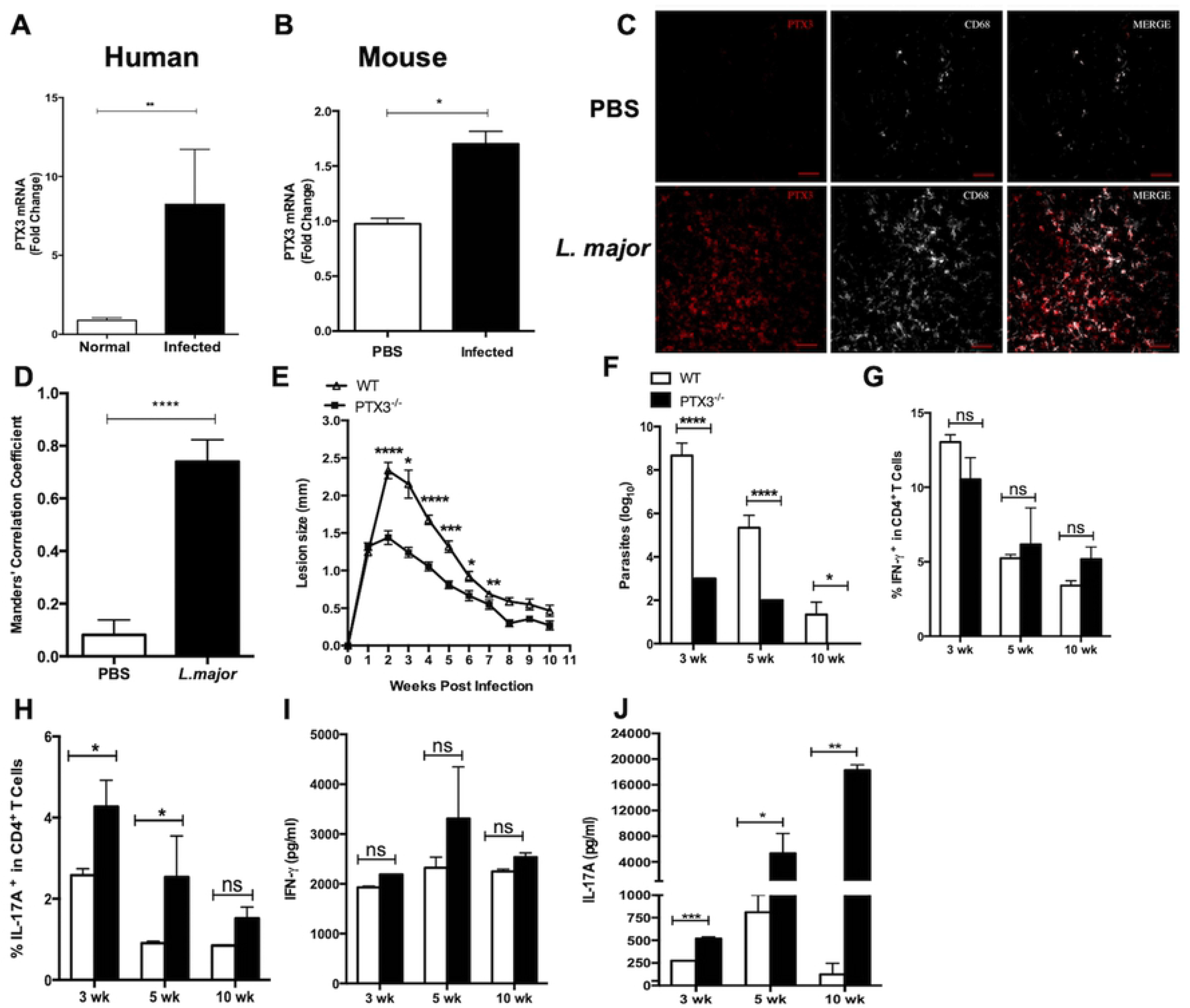
PTX3 expression is increased in CL lesion and mediates susceptibility to *L. major* infection. Skin biopsies from normal (n = 3) and patients with CL (n = 17) were assessed for expression of PTX3 mRNA by RT-PCR (A). Wild type (WT) C57BL/6 mice (n = 4) were infected in the ear with 2 × 10^6^ stationary-phase *L. major* promastigotes and after 7 days, the expression of PTX3 mRNA at the infection site was assessed by RT-PCR using PBS-treated contralateral ears as controls (B). Representative confocal micrographs of ear sections after PBS or *L. major* injection. 10 µm sections of infected or control ears were stained for PTX3 (red) and CD68 (white) and visualized by confocal microscopy (C). Scale bar = 20 µm. Manders’ Correlation Coefficient (MCC) of PTX3 signals in CD68^+^ cells (D). WT and PTX3 deficient (PTX3^-/-^) mice were infected with *L. major* in the right hind footpad and lesion size was measured weekly with digital calipers (E). At the indicated times, infected mice (n = 5 mice per each time point) were sacrificed and parasite burden in the infected footpads was determined by limiting dilution (F). At sacrifice, single cell suspensions of the spleens from infected WT and PTX3^-/-^ mice were analyzed directly *ex vivo* by flow cytometry for the frequency of IFN-γ (G) and IL-17A (H) -producing CD4^+^ T cells. The spleen cells were also stimulated with SLA (50 μg/ml) for 72 h and the levels of IFN-γ (I) and IL-17A (J) in the cell culture supernatant fluids were determined by ELISA. Results are representative of two (B and C) or three (D-J) independent experiments with similar results. *, *p* < 0.05; **, *p* < 0.01; ***, *p* < 0.005; ****, *p* < 0.0001.

The increased expression of PTX3 at lesion sites during *Leishmania* infection suggests that it could mediate either host susceptibility or resistance. To determine this, we compared the outcome of *L. major* infection in wild type (WT) and PTX3 deficient (PTX3^-/-^) mice. At different times after infection, PTX3^-/-^ mice had significantly (p < 0.01-0.0001) smaller lesion size compared to their WT counterparts (Fig 1E). This smaller lesion size corresponded with significantly (p < 0.01-0.0001) lower parasite burden in PTX3^-/-^ mice at 3, 5 and 10 weeks post-infection compared to their WT counterpart mice (Fig 1F). These results indicate that PTX3 negatively regulate disease pathogenesis and possibly immunity during CL.

### Enhanced resistance in PTX3 deficient mice is associated with increased IL-17A production

Next, we assessed cytokine production in the spleen and lymph nodes draining (dLN) the infection site, since resistance to CL is usually associated with robust IFN-γ and reduced IL-10 production by CD4^+^ T cells. Surprisingly, we observed comparable frequencies of IFN-γ^+^CD4^+^ T cells (Fig 1G, Supplementary Fig 2) and IL-10^+^CD4^+^ T cells (Supplementary Fig 3) in both the spleen and dLNs from infected WT and PTX3^-/-^ mice when assessed directly *ex vivo*. Interestingly, we found that PTX3^-/-^ mice had significantly (p < 0.05) higher frequencies of IL-17A^+^CD4^+^ T cells at 3, 5 and 10 weeks post-infection compared to their WT counterparts (Fig 1H). This was consistent with significantly elevated levels of IL-17A (Fig 1J) in the culture supernatant fluids of soluble *Leishmania* antigen (SLA)-stimulated splenocytes isolated from PTX3^-/-^ mice. In contrast and consistent with the flow cytometry data, the levels of IFN-γ (Fig 1I) and IL-10 (Supplementary Fig 3) in the culture supernatant fluids were comparable between WT and PTX3^-/-^ mice. Collectively, these results indicated that the enhanced resistance to *L. major* infection in PTX3^-/-^ mice was not due to enhanced Th1 response, but rather due to stronger Th17 responses in the absence of PTX3 signaling.

### PTX3 deficiency enhances Th17 but does not affect Th1 polarization *in vitro*

Given that we found enhanced IL-17 production following infection of PTX3^-/-^ mice and virtually undetectable levels of IL-17 mRNA in skin biopsies from severe DCL patients (Supplementary Fig 1B), we speculated that PTX3 might negatively regulate Th17 response. To test this, we performed *in vitro* Th17 and Th1 polarization studies using splenocytes obtained from WT and PTX3^-/-^ mice. Data presented in Figs 2A & C show significantly (p < 0.03) higher frequencies of CFSE^lo^CD4^+^IL-17A^+^ T cells in PTX3^-/-^ splenocytes compared to their WT counterparts. In contrast and consistent with our mouse infection studies, the frequency of CFSE^lo^ CD4^+^IFN-γ^+^ T cells in both WT and PTX3^-/-^ splenocytes under Th1 polarizing conditions were comparable (Figs 2B and E). We confirmed these findings by ELISA, which showed higher levels of IL-17A in cell culture supernatants of PTX3^-/-^ splenocytes under Th17 polarization condition (Fig 2D) but comparable levels of IFN-γ in both WT and PTX3^-/-^ splenocytes under Th1 polarization condition (Fig 2E). We also observed similar increased frequencies of Th17 cells using purified CD4^+^ T cells from PTX3^-/-^ spleens under Th17 polarizing conditions (Supplementary Fig 4). These results show that deficiency of PTX3 potentiates Th17 differentiation and IL-17A production.

**Fig 2:**
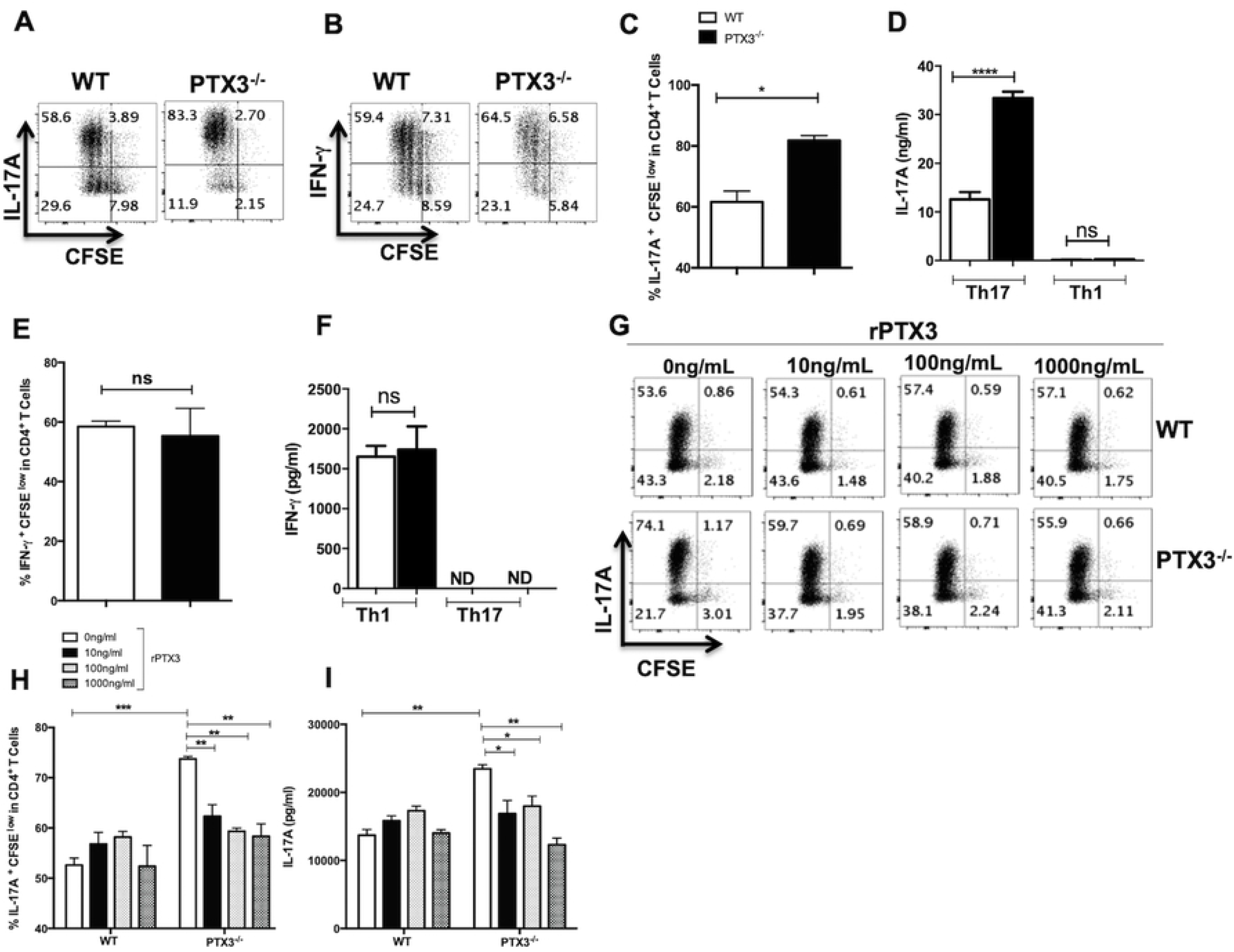
Deficiency of PTX3 enhances Th17 differentiation *in vitro*. Splenocytes from WT and PTX3^-/-^ mice were labeled with CFSE dye and stimulated *in vitro* with soluble anti-CD3 and anti-CD28 antibodies under Th1 or Th17 polarizing conditions. After 72 h, the cells were stained for IFN-γ and IL-17A and the frequencies of IFN-γ− and IL-17A-secreting CD4^+^ T cells were determined by flow cytometry. Shown are representative dot plots (A and B) and bar graphs (C and E) showing the percentage of CFSE^lo^IL-17A^+^ (A and C) and CFSE^lo^IFN-γ^+^ (B and E) CD4^+^ T cells. The levels of IL-17A (D) and IFN-γ (F) in the culture supernatant were assayed by ELISA. CFSE labeled splenocytes from WT and PTX3^-/-^ mice were stimulated with soluble anti-CD3 and anti-CD28 antibodies under Th17 polarizing conditions in presence or absence of different concentrations of rPTX3. After 72 h, the frequency of IL-17A-secreting CD4^+^ cells was determined by flow cytometry. Representative dot plots (G) and bar graphs (H) show the percentage of CFSE^lo^CD4^+^IL-17A^+^ T cells under various conditions. The levels of IL-17A in the culture supernatant were assayed by ELISA (I). Results are representative of three independent experiments with similar results. *, *p* < 0.05; ****, *p* < 0.0001.

The preceding findings suggest that PTX3 may be a negative regulator of Th17 response. To directly test this, we added recombinant PTX3 (rPTX3) to cultures of WT and PTX3^-/-^ splenocytes under Th17 polarization condition. We observed that addition of rPTX3 significantly reduced the frequency of Th17 cells and the production of IL-17A by splenocytes (Figs 2G-I) or purified CD4^+^ T cells (Supplementary Fig 4) from PTX3^-/-^mice. Collectively, these findings directly confirm that PTX3 is a negative regulator of Th17 differentiation and IL-17 production by CD4^+^ T cells.

### PTX3 negatively regulates Th17 specific transcription factors

The observation that rPTX3 suppressed polarization of purified CD4^+^ T cells from PTX3^-/-^ mice into Th17 cells suggests that it may directly affect crucial transcription factors involved in the differentiation of CD4^+^ T cells into Th17 cells. Therefore, we performed RT-PCR to determine mRNA levels of key Th17 transcription factors in WT and PTX3^-/-^ splenocytes under Th17 polarizing conditions in the presence or absence of rPTX3. As expected, there was significantly increased mRNA expression of IL-17A in total splenocytes (Fig 3A) or purified CD4^+^ (Supplementary Fig 5) from PTX3^-/-^ mice compared to those from their WT counterpart mice and this was inhibited by addition or rPTX3. Concomitantly, there was approximately 2-4-fold higher expression of RORγt, STAT3, IRF4, BATF and AhR mRNA in Th17 polarized whole splenocytes (Figs 3B-F) or purified CD4^+^ T cells (Supplementary Fig 5) from PTX3^-/-^ mice compared to those from WT mice. Addition of rPTX3 to PTX3^-/-^ splenocytes resulted in significant inhibition of RORγt and AhR expression in comparison to untreated controls (Figs 3B & F). Similar reduction in the levels of STAT3 was also observed in rPTX3 treated PTX3^-/-^ Th17 cells although these were not statistically significant (Figs 3C & D). Taken together, these findings show that PTX3 negatively regulates Th17 responses by downregulating the expression of Th17 specific transcription factors.

**Fig 3:**
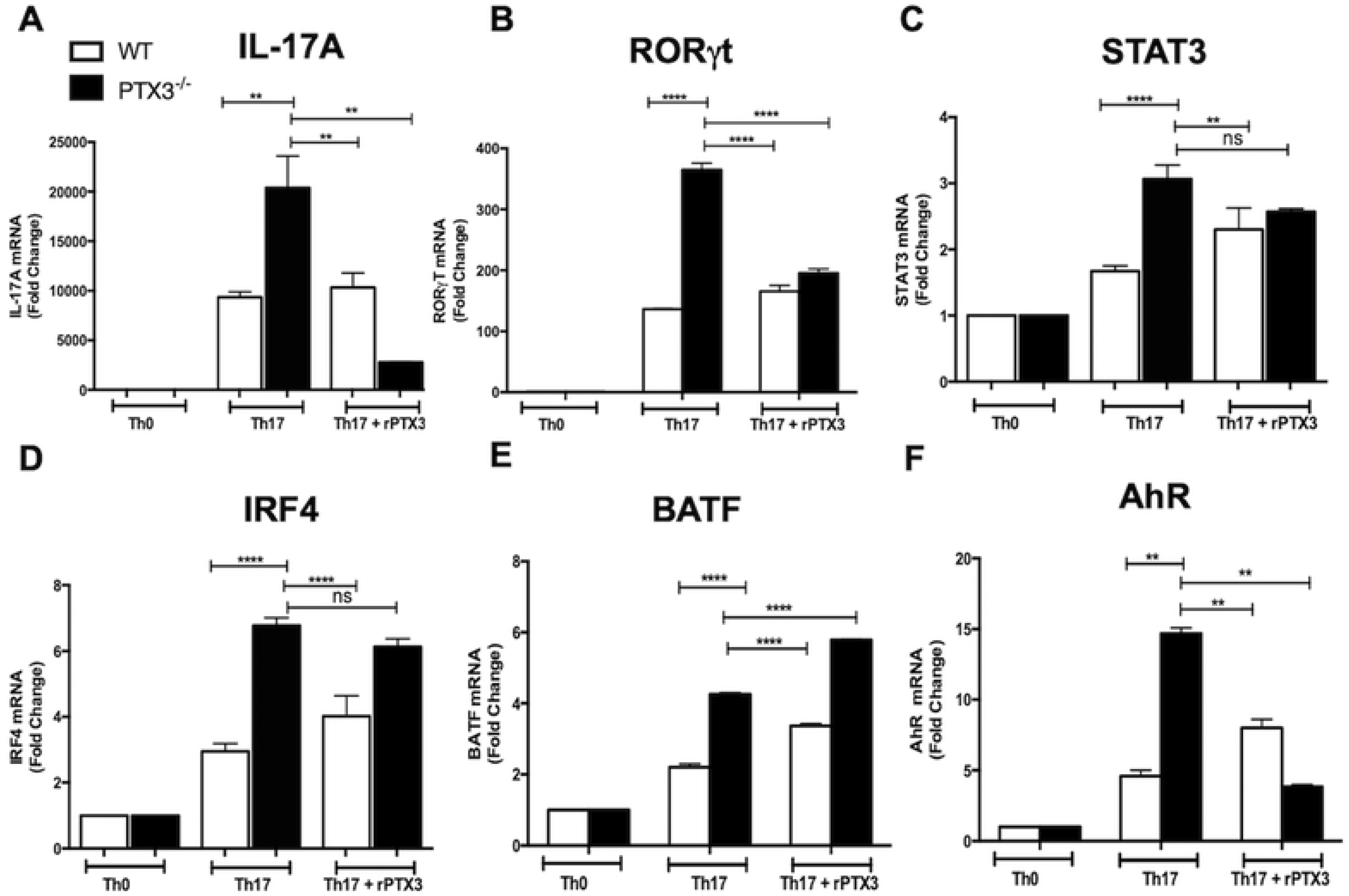
PTX3 negatively regulates Th17-specific transcription. Splenocytes from WT and PTX3^-/-^ mice were stimulated with soluble anti-CD3 and anti-CD28 antibodies under Th17 polarizing conditions in presence or absence of rPTX3 (200 ng/mL). After 72 h, total RNA was isolated and mRNA levels of IL-17A (A), RORγt (B), STAT3 (C), IRF4 (D), BATF (E) and AhR (F) were determined by RT-PCR. Results are representative of three independent experiments with similar results. **, *p* < 0.01; ***, *p* < 0.005; ****, *p* < 0.0001

### Dendritic cells from PTX3^-/-^ mice produce more IL-6 and contribute to increased Th17 responses

Dendritic cells (DCs) present pathogen-derived antigenic peptides to naïve CD4^+^ T cells to initiate antigen-specific T-helper cell activation and differentiation towards specific effector subsets (38-42). Because we found that the absence of PTX3 augmented Th17 responses, we assessed whether deficiency of PTX3 affected DC responses that could favor Th17 differentiation. Splenic CD11c^+^ cells from PTX3^-/-^ mice produced higher amounts of IL-6 (Fig 4A) and IL-12p40 (Fig 4B) compared to those from WT mice following LPS stimulation. Similarly, the expression of IL-6 by MHC-II^+^ CD11c^+^ cells (Figs 4D & E) at the cutaneous site of *L. major* infection was significantly (p < 0.05) higher in infected PTX3^-/-^ mice compared to their WT counterpart controls. These observations suggest that deficiency of PTX3 in DCs leads to enhanced levels of IL-6 during *L. major* infection, which could further contribute to increased Th17 responses in these mice. To confirm this, we co-cultured DCs from WT and PTX3^-/-^ mice with *Leishmania*-PEPCK TCR-transgenic CD4^+^ T cells (1:10) in presence of PECPK peptide. We observed higher frequencies of CD4^+^IL-17A^+^ T cells in co-cultures of PTX3^-/-^ DCs and PEPCK TCR-transgenic CD4^+^ T cells compared to those of WT DCs (Figs 4F and G). Collectively, these findings confirm that PTX3^-/-^ DCs are capable of augmenting Th17 responses during *L. major* infection possibly via regulating IL-6 levels.

**Fig 4:**
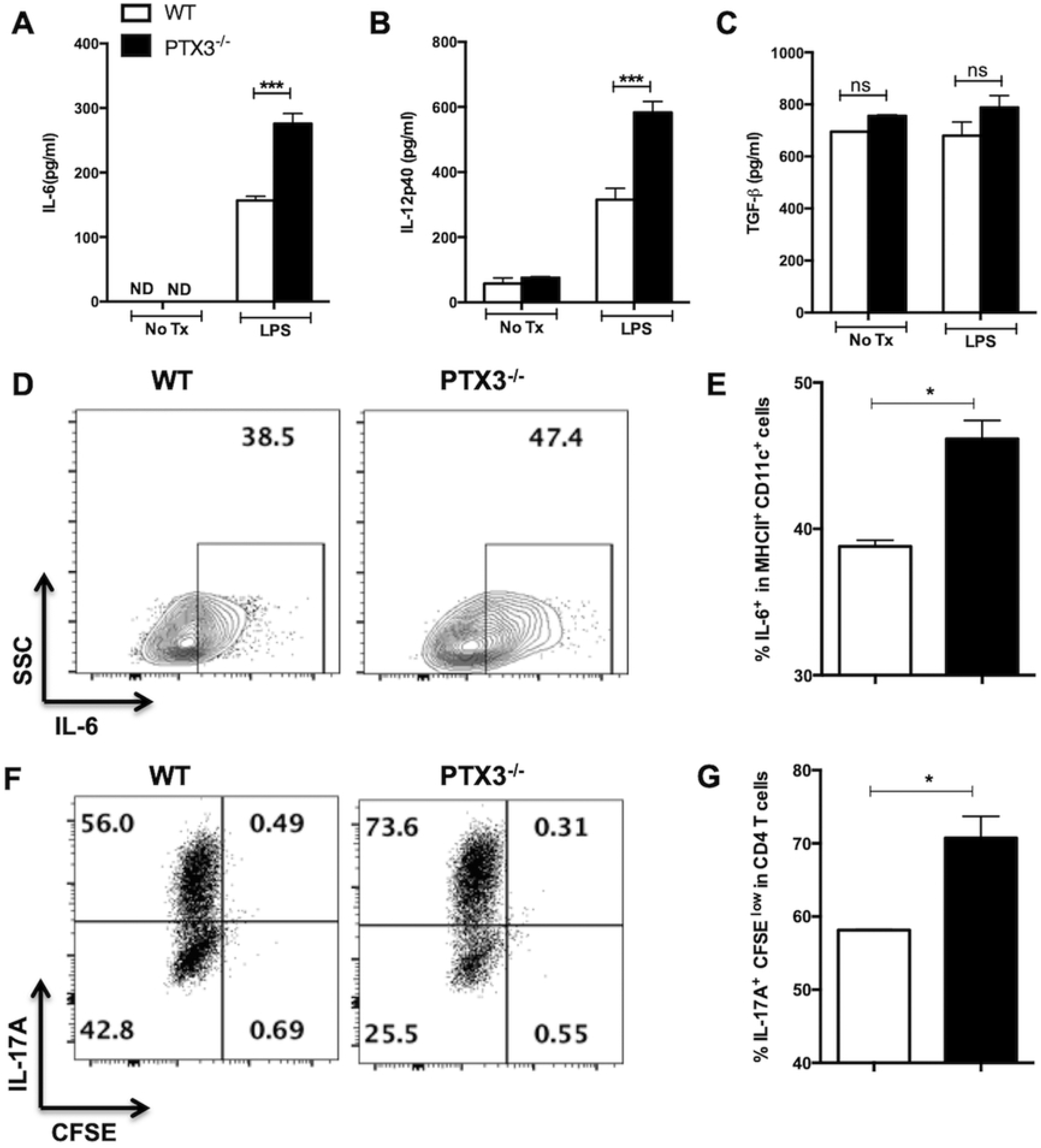
PTX3 regulates DC IL-12 and IL-6 production and function. Highly purified splenic CD11c^+^ cells from naïve WT and PTX3^-/-^ mice were either unstimulated (NoTx) or stimulated with LPS (1 μg/mL) and after 24 h, the cell culture supernatant fluids were collected and assayed for IL-6 (A), IL-12p40 (B) and TGF-β (C) by sandwich ELISA. WT and PTX3^-/-^ mice were infected with *L. major* and after 4 weeks, spleen cells were assessed for the frequency of IL-6 producing MHCII^+^CD11c^+^ cells by flow cytometry. Shown are representative dot plots (D) and bar graph (E) showing the percentage of MHCII^+^CD11c^+^IL-6^+^ cells. Bone marrow-derived DCs (BMDCs) from WT and PTX3^-/-^ were co-cultured with CFSE labeled PEPCK-specific TCR Tg CD4^+^ T cells at 1:10 ratio in presence of PEPCK peptide. After 72 h, the frequency of CD4^+^IL-17A^+^ T cells was determined by flow cytometry. Shown are representative dot plots (F) and bar graph (G) showing the percentage of CFSE^lo^CD4^+^IL-17A^+^ T cells. Results are representative of two independent experiments with similar results. *, *p* < 0.05; ***, *p* < 0.005.

### Enhanced IL-17A responses contribute to increased resistance of PTX3 deficient mice to *L. major* infection

Although some reports have suggested that IL-17 plays a pathogenic role in leishmaniasis (11-13), others showed that they play a protective role (7, 9, 10). Because we found that enhanced resistance of PTX3^-/-^ mice to *L. major* was not associated with superior IFN-γ response, we postulated that the enhanced resistance was mediated by increased IL-17 response. To determine this, we performed an *in vivo* neutralization of IL-17A in *L. major*-infected WT and PTX3^-/-^ mice and monitored lesion size and parasite burden at 4 weeks post-infection. IL-17A neutralization in infected PTX3^-/-^ mice resulted in increased lesion size (Fig 5A) and a concomitant increase in parasite burden (Fig 5B) compared with untreated PTX3^-/-^ mice.

**Fig 5:**
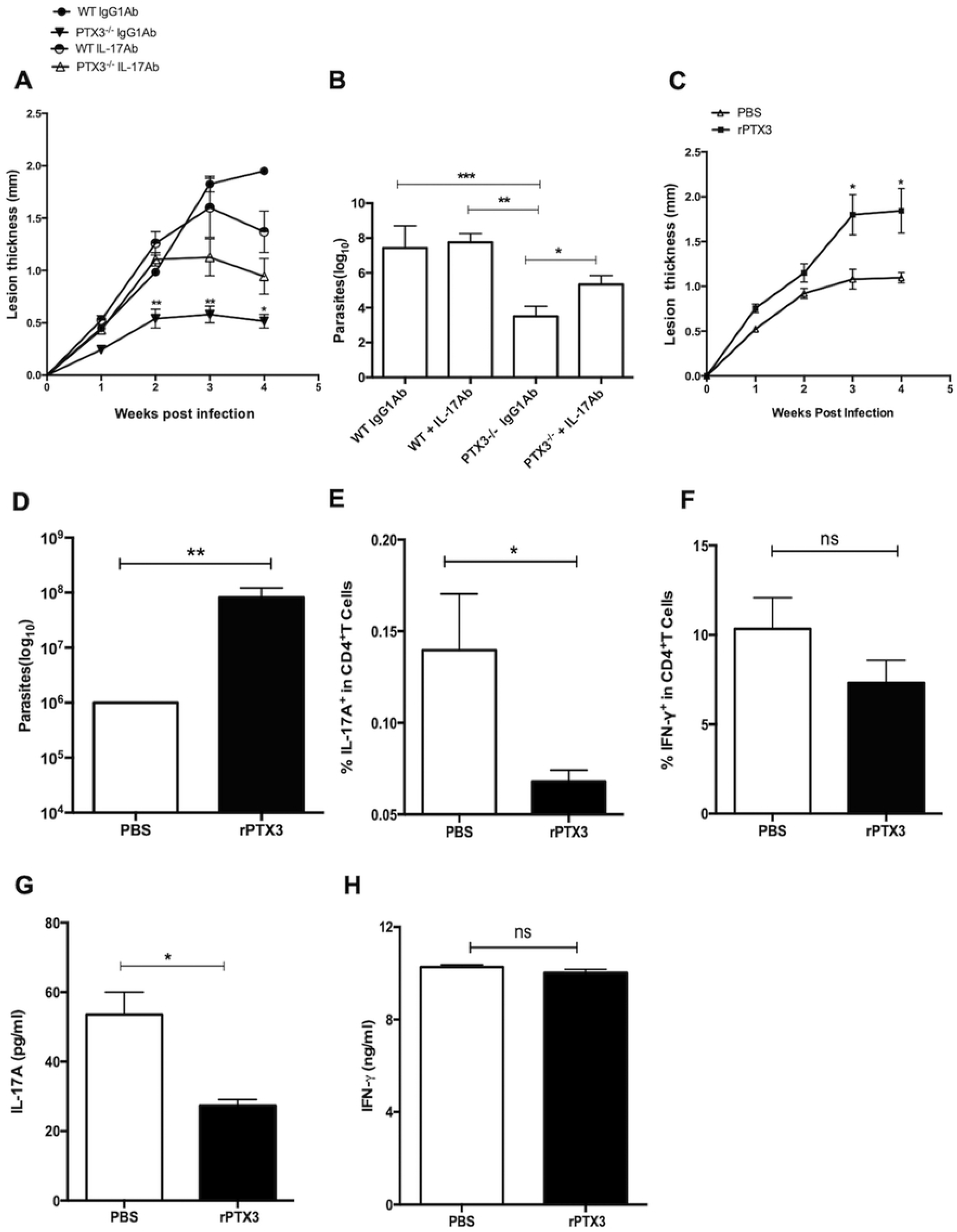
*In vivo* neutralization of IL-17A or rPTX3 treatment increase susceptibility to *L. major* infection. WT and PTX3^-/-^ mice (n = 5 per group) were treated (i.p) with either control Ig or anti-IL-17A neutralizing antibody one day before *L. major* infection and continued once weekly for additional 4 weeks. Lesion size (A) was measured weekly and mice were sacrificed after 4 weeks to estimate parasite burden in the infected footpads (B). Groups of WT mice were infected with *L. major* and one day post-infection, mice were divided into two groups (6 mice per group) and treated intralesionally with either PBS or rPTX3 (0.5 μg/mouse) 3 times weekly. Lesion size (C) was measured weekly and after 4 weeks, mice were sacrificed and parasite burden in the infected footpads was determined by limiting dilution (D). Cells from the dLNs were assessed directly *ex vivo* for the frequency of IL-17-(E) and IFN-γ (F) producing CD4^+^ T cells by flow cytometry. Some dLN cells were stimulated with SLA (50 μg/ml) for 72 h and the levels of IL-17A (G) and IFN-γ (H) in the cell culture supernatant fluids were determined by ELISA. Results are representative of two independent experiments with similar results. *, *p* < 0.05; **, *p* < 0.01; ***, *p < 0.001*.

Next, we evaluated if administration of rPTX3 to WT mice could lead to increased susceptibility to *L. major* infection. We infected WT mice with *L. major* and administered rPTX3 intralesionally once a week for 3 weeks. WT mice treated with rPTX3 had increased lesion size (Fig 5C) that corresponded with significantly increased parasite burden (Fig 5D) compared to PBS treated controls. The enhanced susceptibility following rPTX3 treatment was accompanied by significant (p < 0.05) reduction in the frequency of CD4^+^IL-17A^+^ T cells in the dLNs and spleen compared to PBS treated controls (Fig 5E and Supplementary Fig 6). Consistent with previous findings (Figs 2B & F), there was no difference in the frequency of CD4^+^IFN-γ^+^ T cells in dLNs and spleen of both rPTX3 and PBS treated groups (Fig 5F and Supplementary Fig 6). We confirmed the above results by ELISA, which showed increased levels of IL-17A (Fig 5G and Supplementary Fig 6) and unchanged levels of IFN-γ (Fig 5H and Supplementary Fig 6) in cell culture supernatant fluids of SLA-stimulated dLN and spleen cells from rPTX3-treated mice. Collectively, these findings confirm that PTX3 enhances susceptibility to *L. major* infection by downregulating IL-17A response.

### IL-17A synergizes with IFN-γ to mediate effective parasite killing in macrophages

*Leishmania* resides inside host macrophages and their clearance requires activation of infected cells by IFN-γ leading to the production of reactive oxygen and nitrogen intermediates (43). To fully understand how deficiency of PTX3 enhances resistance to *L. major* infection, we compared the uptake, replication and killing of parasites in macrophages, from WT and PTX3^-/-^ mice. Both WT and PTX3^-/-^ macrophages had similar parasite uptake as seen by equivalent numbers of amastigotes in WT and PTX3^-/-^ cells at 6 h post-infection (Figs 6A and B). In addition, both WT and PTX3^-/-^ macrophages had similar number of amastigotes at 24, 48 and 72 h post-infection (Figs 6A and B), suggesting that deficiency of PTX3 had no effect on parasite replication in infected cells. Furthermore, both infected WT and PTX3^-/-^ macrophages had comparable ability to kill parasites following activation with LPS or IFN-γ (Fig 6C).

**Fig 6:**
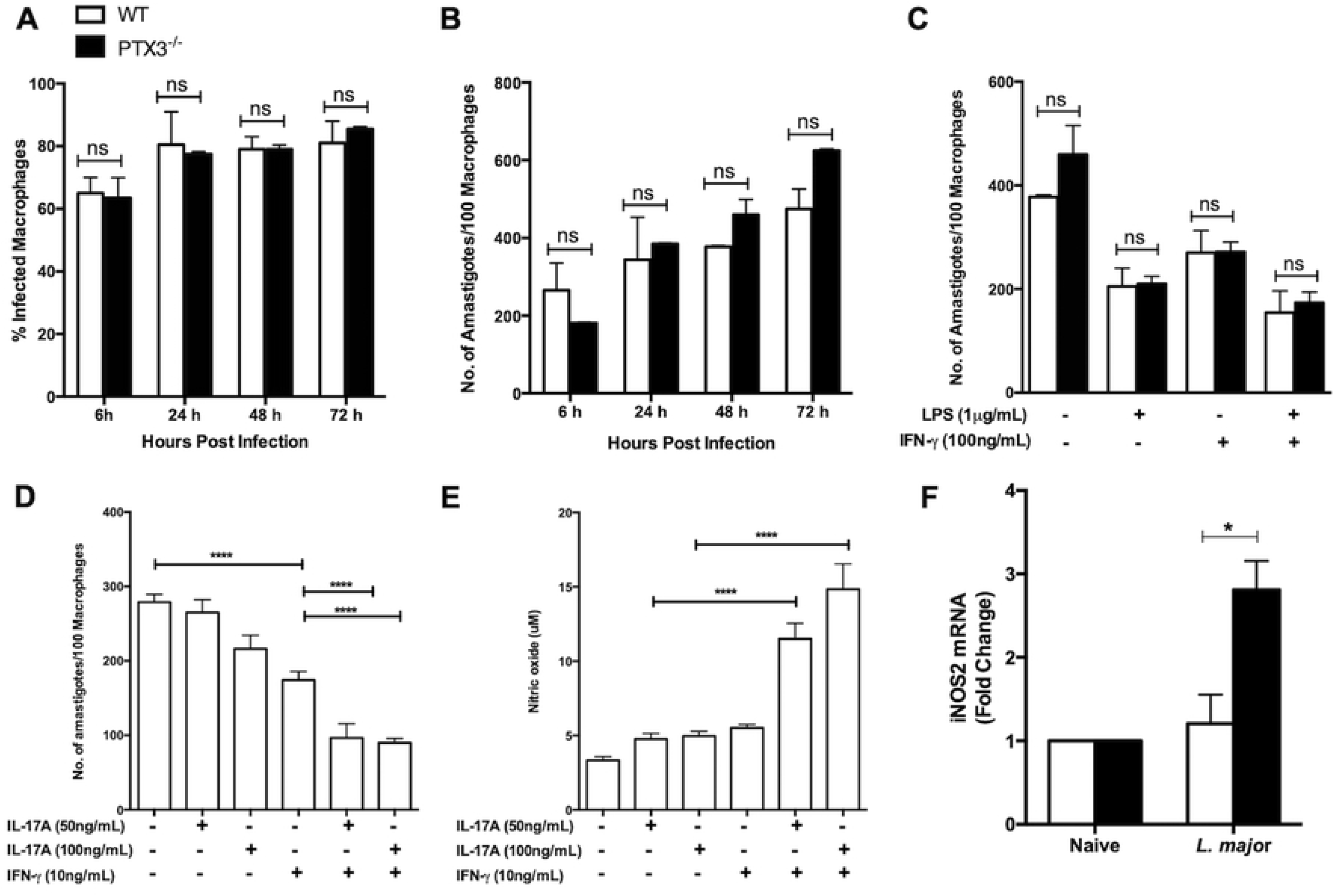
PTX3 synergizes with IFN-γ to mediate killing of *L. major* in infected macrophages. Bone marrow-derived macrophages from WT and PTX3^-/-^ were infected with *L. major* and at indicated times, cytospin preparations of infected cells were stained with Wright– Giemsa stain and the percent infection (A) and total number of parasites per 100 cells (B) were determined by microscopy. In another independent experiment, infected bone marrow-derived macrophages from WT and PTX3^-/-^ were stimulated with either LPS (1μg/mL), IFN-γ (100 ng/mL) or both for 48 h and the number of parasites per 100 cells was determined (C). Bone marrow-derived macrophages from PTX3^-/-^ were infected with *L. major* in presence or absence of IL-17A (50 & 100 ng/mL) or IFN--γ (10 ng/mL) alone or both. After 48 h, cytospin preparations were stained with Wright–Giemsa stain and the number of parasites per 100 cells was determined by microscopy (D) and the concentration of nitrite in the supernatant was determined (E). WT mice (n = 4) were infected with *L. major* promastigotes and after 4 weeks, the expression of iNOS2 mRNA at the infection site was assessed by RT-PCR using PBS-treated contralateral footpads as controls (F). Data are shown as mean ± SEM of four to six infection tubes per group and are from single experiment representative of at least three independent experiments. *, *p* < 0.05; ****, *p* < 0.0001.

**Fig 7:**
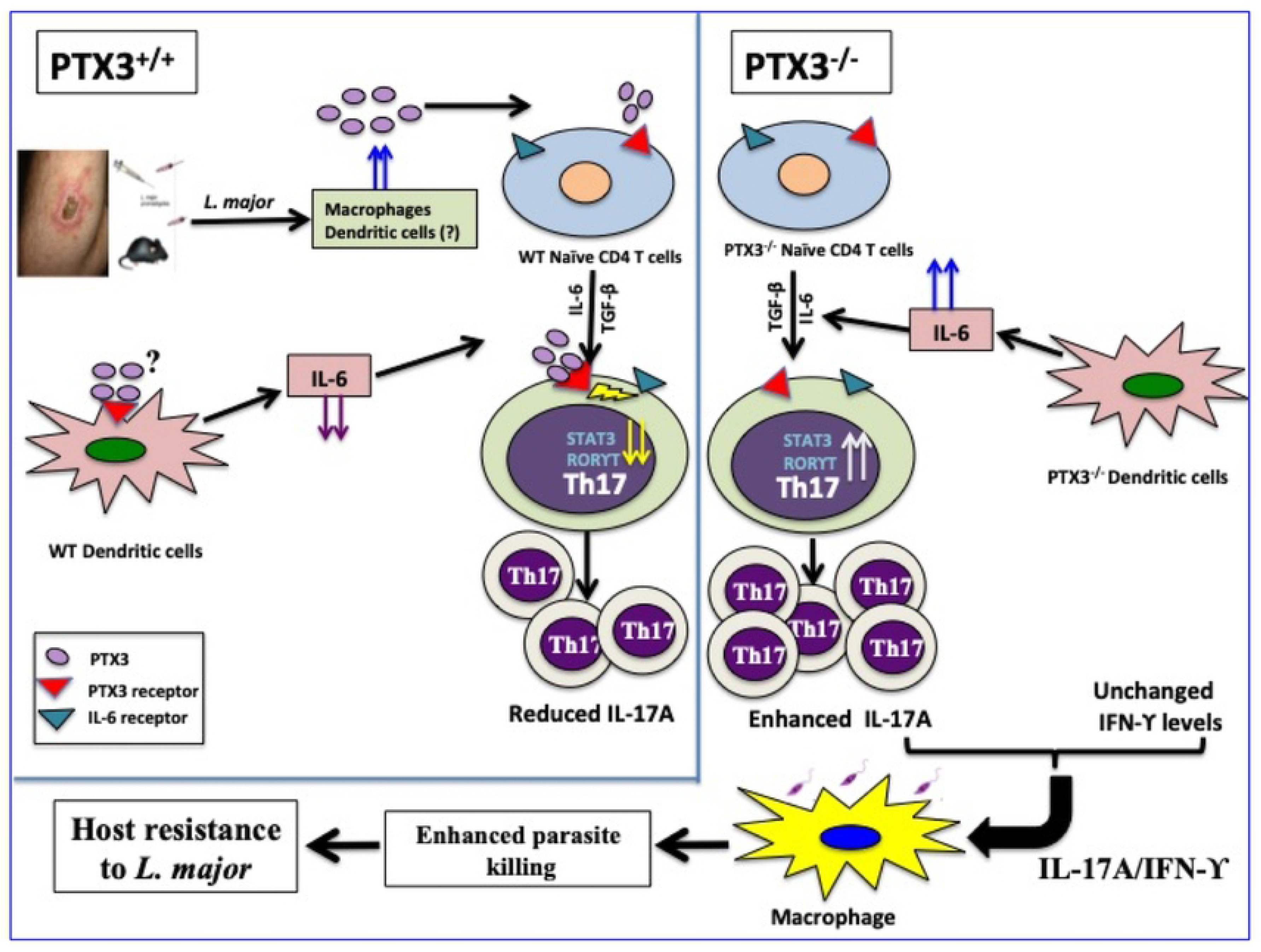
PTX3 mediated regulation of immunity to *Leishmania major*. In a PTX3 competent system, *Leishmania major* infection induces the production of PTX3 from immune cells (such as macrophages and dendritic cells). PTX3 downregulates the production of IL-6 which contributes to suppression of Th17 response. In addition, PTX3 directly inhibits the expression of Th17-specific transcription factors (including RORγT, IRF-4, BATF, AhR and STAT3) further enhancing the suppression of Th17 differentiation and IL-17A production. In the absence of PTX3 (as seen in PTX3 deficient mice), increased expression of IL-6 and Th17-specific transcription factors leads to increased Th17 differentiation and a concomitant increase in the frequency of Th17 cells and increased levels of IL-17A. This enhanced IL-17A production synergizes with IFN-γ to more efficiently activate *Leishmania-*infected macrophages resulting in increased nitric oxide production, enhanced parasite killing and increased host resistance to *L. major*.

Because IL-17 has been proposed to enhance leishmanicidal activity (9) we examined whether the enhanced resistance to *L. major* in PTX3^-/-^ mice was related to IL-17 augmentation of IFN-γ-mediated *Leishmania* killing activity. We primed PTX3^-/-^ macrophages with IL-17 (50 & 100 ng/mL) in the presence or absence of suboptimal dose of IFN-γ (10 ng/mL) and infected them with *L. major*. Results presented as Figs 6D & E show that IL-17 cooperates with suboptimal dose of IFN-γ to mediate increased NO production and more effective parasite killing (compared to treatment with IFN-γ or IL-17 alone). Similarly, we observed increased iNOS2 mRNA expression at the site of infection in PTX3^-/-^ mice compared to WT counterparts (Fig 6F), which confirmed our *in vitro* findings. Collectively, our results show that the enhanced resistance of PTX3^-/-^ mice to *L. major* is due to enhanced IL-17A production, which synergizes with IFN-γ to enhance NO production, leading to more effective killing of parasites in infected macrophages.

## Discussion

Herein we showed that PTX3 levels were elevated in skin lesions from patients and mice suffering from CL, suggesting that this innate pattern recognition molecule may play a critical role in disease pathogenesis. Using a loss of function approach, we showed that deficiency of PTX3 resulted in enhanced resistance to *L. major*, and this was associated with increased IL-17 (but not IFN-γ) response. Neutralization studies showed that the enhanced resistance of PTX3^-/-^ mice to *L. major* is due to enhanced Th17 responses in these mice. In contrast, administration of rPTX3 led to increased susceptibility to *L. major* which was associated with a dramatic downregulation of Th17 responses and IL-17A production by lymph node cells draining the infection site. Using an *in vitro* approach, we showed that CD4^+^ T cells from PTX3^-/-^ mice showed enhanced expression of Th17 transcription factors that drive Th17 differentiation. Collectively, these results show, for the first time, that PTX3 is a negative regulator of Th17 response during *Leishmania major* infection.

The expression of PTX3, a key molecule of the innate immune defense system, is upregulated in response to different stimuli such as inflammatory cytokines (IL-1β, TNFα), TLR agonists (e.g. LPS); distinct microbial associated molecular patterns (such as OmpA, lipoarabinomannans) and some pathogens (*E. coli*, *S. aureus*) (19, 44-46). Studies with PTX3^-/-^ and PTX3-overexpressing mice have shown that PTX3 mediates protective immunity to various pathogens including influenza virus, *Aspergillus fumigatus*, and *Pseudomonas aeruginosa* (15, 33). While correlative studies in human leishmaniasis patients suggest that PTX3 may play a key role in disease pathogenesis, no study has directly demonstrated this and/or showed the mechanism through which this would occur. Results from our studies clearly show that *Leishmania* infection induces PTX3 expression and this blocks effective parasite control by suppressing protective Th17 and IL-17A responses.

IL-17A is a proinflammatory cytokine produced primarily by CD4^+^ Th17 cells although other cell types, such as CD8^+^ T cells, γδ T cells, invariant natural killer T (iNKT) cells and neutrophils are known to also secrete it. Binding of IL-17A to the IL-17 receptor, which is expressed on many cells including macrophages, initiates a strong signaling cascade that leads to expression of inducible nitric oxide synthase, GMCSF, proinflammatory cytokines, antimicrobial peptides and chemokines (47, 48) that are important for host protection from many pathogens such as bacteria (49), fungi (50) and trypanosomes (51). We showed the PTX3 negatively regulates IL-17A production during *L. major* infection since its deficiency led to increased frequency of IL-17A producing CD4^+^ T cells in the dLNs and spleens of infected mice. In support of this, we found that the increased expression of PTX3 mRNA in biopsy samples from DCL patients was associated with suppression of IL-17A mRNA in these tissues. Indeed, neutralization of IL-17A abolished the enhanced resistance of infected PTX3^-/-^ mice while rIL-17A treatment conferred enhanced resistance to WT mice as evidenced by significantly reduced lesion size and parasite burden. Furthermore, we found that IL-17A synergizes with IFN-γ to mediate enhanced NO production and a concomitant more efficient parasite killing in infected macrophages.

The role of IL-17 in the pathogenesis of CL is controversial. While a report suggests that IL-17A mediates susceptibility to *L. major* infection in mice by regulating CXCL2 levels and neutrophil migration to the site of infection (12), observations in human patients suggest that its increased expression may contribute to better disease outcome. Increased IL-17A levels has been shown to correlate with better disease outcome in subclinical *L. braziliensis* patients (52). Similarly, studies have shown that IL-17A (possibly derived from Th17 cells) mediates protective immunity in patients against *L. infantum, L. major* and human post kala-azar dermal leishmaniasis (9, 10, 53). In line with this, we found that the expression of IL-17A mRNA was highly suppressed in tissue biopsies from DCL patients. Importantly, the pathways leading to IL-17A production in infected mice and patients are not known. In the present study, we showed that PTX3 is a key molecule that regulates IL-17A response in CL. The expression of PTX3 in cutaneous lesions was directly correlated with clinical pathology of the disease such that PTX3 levels were highest in patients displaying disseminated (54, 55) and recidivous lesions (56). In contrast, PTX3 levels were inversely correlated with the level of IL-17A, such that individuals exhibiting disseminated disease had undetectable levels of IL-17 mRNA in their lesions. Treatment of mice with rPTX3 resulted in enhanced susceptibility to *L. major* infection due to suppressed Th17 responses. These findings are consistent with previous studies on *Aspergillus* where rPTX3 treatment led to suppression of Th17 responses (57).

Th17 cell differentiation is driven by TGF-β and IL-6 (58, 59) and regulated by some key transcription factors including RORγt (59), STAT3 (60), IRF4 (61), BATF (62) and AhR (63). In the presence of IL-6 and TGF-β, the cooperative binding of BATF, IRF4 and STAT3 with AhR contributes to initial chromatin accessibility and subsequent recruitment of RORγt to regulate activation of Th17-relevant genes (64, 65). We observed increased expression of RORγt, STAT3, IRF4, BATF and AhR in CD4^+^ T cells from PTX3^-/-^ under Th17 polarizing conditions which correlated with enhanced Th17 and IL-17A responses. Addition of rPTX3 significantly suppressed the expression of RORγt, STAT3 and AhR, transcription factors and IL-17A production. These findings show that PTX3 is capable of downregulating these multiple transcription factors to suppress Th17 response. Our findings are in line with a previous study which showed that deficiency of PTX3 led to enhanced Th17 response via upregulation of STAT3 in murine model of allergic asthma (14).

Although different inflammatory cytokines (including IL-1β, TNFα), TLR agonists (e.g. LPS); PAMPS and pathogens have been shown to induce PTX3 expression in cells (19, 44-46), PTX3 induction by *Leishmania* has never been reported. We found that *Leishmania* infection induced massive expression of PTX3 by CD68^+^ cells at the site of infection. Whether this induction is direct (via parasite-derived molecules) or indirect (via production of cytokines by infected cells) remains unknown. It is conceivable that some parasite-derived molecules such as LPG or GP63 could play a role in the induction of PTX3 following infection in order to downregulate protective Th17 response. This could be a novel evasion strategy employed by *Leishmania* to subvert the host immune response thereby favoring its survival within the infected host.

Taken together, the present study reveals a hitherto unknown role of PTX3 in regulating host immunity by suppressing Th17 and IL-17A responses. Following *L. major* infection, the production of PTX3 in WT mice limits protective IL-17 response by downregulating IL-6 production by DCs and activation of key transcription factors that favor Th17 activation in CD4^+^ T cells. In absence of PTX3 (as seen in PTX3^-/-^ mice), increased production of IL-6 and TGF-β by infected DCs favors optimal differentiation of CD4^+^ T cells into Th17 cells via increased expression of Th17-specific transcription factors like STAT3, AhR RORγT, leading to enhanced production of IL-17A. Our studies clearly highlight the importance of Th17 and IL-17A responses in resistance to CL. The findings that treatment of WT mice with rPTX3 modulated Th17 response without affecting IFN-γ (Th1) response allowed us to directly demonstrate the contribution of IL-17 in resistance to CL. The findings clearly establish IL-17A as a critical cytokine that contributes to optimum resistance to CL. They show that IL-17A synergizes with IFN-γ to more efficiently activate infected macrophages leading to increased production of NO that mediates effective parasite killing. These findings suggest that PTX3 could be a therapeutic target for regulating immunity to CL.

## Materials and Methods

### Mice

Heterozygous female PTX3^-/+^ and homozygous male PTX3^-/-^ (129SvEv/Bl/6 background) mice were bred at the University of Manitoba Central Animal Care Services breeding facility. Homozygous female PTX3^-/-^ and their female homozygous wild type littermate mice (6-8 weeks old) were used in in the studies. Additionally, in some studies female PEPCK TCR-transgenic on a C57BL/6 genetic that recognizes *Leishmania* specific PEPCK peptide, were used.

### Human CL patients

This study was conducted in Jequiriça, Bahia, Brazil, a well-known area of *L. braziliensis* transmission. Participants included 3 healthy endemic controls and 17 patients with Tegumentary leishmaniasis with cutaneous lesions typical of *Leishmania* infection and a positive Montenegro skin test (Supplementary Table 1).

### Ethics Statement

All mice were kept at the University of Manitoba Central Animal Care Services (CACS) facility in accordance to the Canadian Council for Animal Care guidelines. The University of Manitoba Animal Use Ethics Committee approved all studies involving animals, including infection, humane endpoints, euthanasia and collection of samples(Protocol Numbers 17-007, AC 11232).

Research on human CL patients was conducted with the approval of the Ethical Committee of Hospital Santa Izabel-Santa Casa de Misericórdia da Bahia (Salvador, Bahia, Brazil; 1.163.870) and Comissão Nacional de Ética em Pesquisa (CEP, Brazilian National Ethics Committee, Brazil). Informed consent was obtained from each participant. All methods were performed in accordance with the guidelines and regulations determined by CEP. All human subjects that were part of this study were adults, and informed written consent was obtained from them.

### Parasites and infection

*L. major* parasites [MHOM/IL/80/Friedlin (FN)] were cultured at 26°C in M199 medium (HyClone, Logan, UT) supplemented with 20% heat-inactivated FBS (HyClone), 2 mM L-glutamine, 100 U/mL penicillin, and 100 μg/mL streptomycin (Invitrogen Life Technologies, Burlington, Ontario, Canada). For infection, mice were injected in the right hind footpad with 2 × 10^6^ stationary phase promastigotes in 50 μL PBS as previously described (66). In some experiments, the mice were injected in the right ear lobe with 2 × 10^6^ stationary phase promastigotes in 10 μL PBS. Lesion sizes were monitored weekly by measuring footpad swellings with digital calipers. Parasite burden in the infected footpads was determined by limiting dilution assay.

### *In vivo* blockade of IL-17A, rPTX3 treatment and estimation of parasite burden

For *in vivo* neutralization of IL-17A, WT and PTX3^-/-^ mice were injected with anti–IL-17A (clone 17F3) mAb or control Ig (1 mg/mouse) i.p. 1 day before infection with *L. major*. Antibody treatment was continued once weekly at 0.5 mg/mouse for additional 4 weeks. The lesion thickness was monitored weekly and mice were sacrificed after 4 weeks post infection to determine parasite burden. To assess the impact of PTX3 on disease outcome, infected WT mice were either injected locally (intraleisonally) with PBS or rPTX3 (0.5 mg in 50 ul PBS) thrice weekly and lesion thickness was monitored weekly. Treated mice were sacrificed at 4 weeks post-infection to determine immune response in spleens and dLNs and parasite burden in the infected footpads. Parasite burden in the infected footpads was quantified by limiting dilution analysis as previously described (67).

### *In vitro* recall response and intracellular cytokine staining

At various times post-infection, infected mice were sacrificed and the draining popliteal lymph nodes or cervical lymph nodes (ear infection) were harvested and made into single-cell suspensions. Cells were washed, resuspended at 4 million/ml in complete medium (DMEM supplemented with 10% heat-inactivated FBS, 2 mM glutamine, 100 U/ml penicillin, and 100 μg/ml streptomycin), and plated at 1 ml/well in 24-well tissue culture plat (Falcon, VWR Edmonton, AB, Canada). Cells were stimulated with SLA (50 μg/ml) for 72 h, and the supernatant fluids were collected and stored at −20°C until assayed for cytokines by ELISA.

### Cytokine ELISAs and NO

IL-17A IFN-γ and IL-10 concentrations in cell culture supernatant fluids were measured by sandwich ELISA using Ab pairs from BD Pharmingen or Biolegend according to manufacturer’s suggested protocols. Nitrite concentrations in BMDM culture supernatants were used as a measure of NO production and quantified using the Griess assay.

### Generation of bone marrow–derived macrophages (BMDM), dendritic cells (BMDC) and *in vitro* infections

Bone marrow–derived DCs (BMDCs) and bone marrow–derived macrophages (BMDMs) were generated from naive WT and PTX3^-/-^ mice as described previously (66). In brief, bone marrow cells were isolated from the femur and tibia of mice and differentiated into macrophages using complete medium supplemented with 30% L929 cell culture supernatant. For BMDC differentiation, the bone marrow cells were grown using complete medium supplemented with 20ng/mL GM-CSF. For infection, BMDMs were incubated with parasites for 6h at a cell/parasite ratio of 1:10 as previously described (66). In some experiments, infected cells were stimulated with IFN-γ (100 and 10 ng/mL), IL-17A (100 and 50 ng/mL) and LPS (1 μg/mL). At different times after infection, parasite numbers inside infected cells were determined by counting Giemsa stained cytospin preparations under light microscope at 100x (oil) objective. In addition, the culture supernatant fluids were also assessed for nitrite concentration. In some experiments, BMDCs were stimulated *in vitro* with LPS (1 μg/mL) for 24 h, and culture supernatant fluids were assayed for TGF-β, IL-12p40 and IL-6 by ELISA.

### Purification of splenic CD4^+^ T cells and CD11c^+^ (dendritic) cells

Splenic CD4^+^ T and CD11c^+^ cells were isolated by negative and positive selection using the StemCell CD4^+^ T and CD11c^+^ cells EasySep isolation kits, respectively, according to the manufacturer’s suggested protocols. The purities of the different cell populations were > 94% (CD4^+^) and 87–93% (CD11c^+^).

### *In vitro* Th1 and Th17 differentiation

Single-cell suspensions from the spleens (whole splenocytes) or highly purified naïve CD4^+^ (CD44^-^CD62^+^) cells from WT and KO mice were labeled with CFSE dye as previously described (68) and cultured in 96-well plates (2 × 10^5^ per well in 200 μL aliquots) in the presence of plate-bound anti-CD3 (1 μg/mL) and anti-CD28 (1 μg/mL) under varying polarizing conditions as follows: Th1, rIL-12 (20 ng/mL) and anti-IL-4 (10 μg/mL); Th17, rTGF-β (10 ng/mL), rIL-6 (100 ng/mL), anti-IL-4 (10 μg/mL), anti-IL-2 (10 μg/mL) anti-IFN-γ (10 μg/mL), and anti-IL-12 (10 μg/mL). All recombinant cytokines were purchased from Peprotech while endotoxin-free mAbs were purchased from BioLegend (San Diego, CA). In some experiments, rPTX3 (R&D system) was added into the cell cultures. After 5 days of culture, the cells were routinely assessed for proliferation and cytokine production by flow cytometry as described below.

### BMDC-T cells Co-culture assays

CFSE labeled highly purified naive CD4^+^ T cells from PEPCK TCR transgenic mice were cultured for 4 days in 96-well plates with LPS-matured BMDCs from WT and PTX3^-/-^ mice at 10:1 (DC to T cell ratio) in presence of PEPCK peptide (5 μM NDAFGVMPPVARLTPEQ, (69) and Th17-polarizing cocktail (as described earlier). After 5 days of culture, the cells were routinely assessed for proliferation and cytokine production by flow cytometry as described below.

### CFSE labeling and proliferation protocol

The CFSE labeling protocol used here has been described previously (68). Briefly, single-cell suspensions from the spleens or dLNs were counted and stained with CFSE dye at 1.25 μM at room temperature in the dark with continuous rocking. After 5 min, staining was quenched with heat-inactivated FBS and the cells were washed, counted, resuspended in complete medium, and used for *in vitro* cultures.

### Quantification of transcript levels by RT-PCR

Total RNA was extracted from murine ear, splenocytes or purified CD4 T cells using the RNeasy Plus Micro Kit. mRNA was reverse transcribed and cDNA was amplified by RT-PCR using SYBR Green chemistry as described previously (70). Murine Primers and reaction conditions were found using the PRIMER BANK website (Massachusetts General Hospital. Primer Bank. http://pga.mgh.harvard.edu/primerbank). Data were normalized to the housekeeping gene β-actin and presented as fold induction over non-polarized splenocytes or CD4^+^ T cells using the delta-delta CT method.

Cryopreserved human skin biopsies from lesions of infected or uninfected people were processed into fine powder using the traditional mortar and pestle system. Total RNA was extracted from these samples using RNeasy mini kit (Qiagen, Venlo, Netherlands) and DNA clean up was performed on-column by DNAse treatment (Qiagen). mRNA was reverse transcribed and cDNA was amplified using Taqman gene expression assays (Applied Biosystems) for PTX3 (Hs00173615_m1), IL17A (Hs00174383_m1) and GAPDH (Hs03929097_g1). All reactions were performed using the standard cycling conditions (Applied Biosystems, Warrington, United Kingdom). Data were normalized to the housekeeping gene GAPDH and presented as fold induction over normal skin (NS) using the delta-delta CT method.

### Flow cytometry analysis

For flow cytometry analysis, splenocytes and dLN cells, were directly surface stained for CD3, CD4, MHC-II and CD11c expression. For intracellular cytokine analysis, surfaced - stained splenocytes and dLN cells were permeabilized with 0.1% saponin (Sigma - Aldrich) in staining buffer and then stained with specific fluorochrome-conjugated mAbs against IL-6, IFN-γ, IL-10, and IL-17 (BioLegend). Samples were acquired on a BD FACSCantor machine and analyzed using FlowJo software (TreeStar Inc, Ashland, OR).

### Confocal microscopy

PBS treated and *L. major* infected ear tissues from C57BL/6 mice were harvested after 3 days post infection. The tissues were fixed for 1h at 4 °C in 4% paraformaldehyde in PBS, and incubated at 4 °C for 1h in 10% and 20% sucrose in PBS, then in 30% sucrose overnight. Tissues were embedded in OCT compound (Fisher Scientific) and cut into 10 μm sections using a cryostat and mounted onto microscope slides. Slides were washed, blocked with Fc blocker (Innovex), 4% mouse serum (ImmunoReagents) and 4% goat serum. The primary antibodies used were rat anti-PTX3 (Enzo Life Sciences) at a 1:400 dilution, and rabbit anti-CD68 (Abcam) at a 1:400 dilution. Secondary antibodies used were AF568-conjugated goat anti-rat (Invitrogen) at 1:1000 dilution, and AF647-conjugated goat anti-rabbit (Invitrogen) at 1:5000 dilution. Slides were stained with Hoechst 33342 (Molecular Probes) for 30 min at 1:2000 dilution and mounted with ProLong Gold (Invitrogen). Images were acquired using the Zeiss AxioObserver confocal microscope. Colocalization analysis (using Manders’ correlation coefficient) was performed using the JACoP plugin in ImageJ.

### Statistics

Results are shown as means ± SE. Results from different groups were compared using Student’s *t*-test or one way Annova. A *p* value of ≤ 0.05 was considered significant.

## Acknowledgments

We thank members of Uzonna lab for their insightful comments.

## Supplementary Figure Legends

**Supplementary Fig 1: Expression of PTX3 and IL-17A in different forms of CL**

Skin biopsies from normal (NS) (n = 3) and patients with different forms of CL: localized CL (LCL, n = 6), *Leishmania* Recidiva Cutis (LRC, n = 5) and Disseminated leishmaniasis (DCL, n = 6) were assessed for expression of PTX3 (A) and IL-17 (B) mRNA by RT-PCR. *, *p* < 0.05.

**Supplementary Fig 2: Absence of PTX3 enhances IL-17A but has no effect on IFN-γ response in *L. major*-infected mice.**

Groups of WT and PTX3^-/-^ mice were with infected with *L. major* and at the indicated times, infected mice (n = 5 per each time point) were sacrificed and single cell suspension of the dLNs were analyzed directly *ex vivo* by flow cytometry for the frequency of IL-17A (A and D) and IFN-γ (B and E) -producing CD4^+^ T cells. Some dLN cells were stimulated with SLA (50 μg/ml) for 72h and the levels of IFN-γ (C) in the cell culture supernatant fluids was measured by ELISA. Results are representative of three independent experiments with similar results. *, *p* < 0.05.

**Supplementary Fig 3: Deficiency of PTX3 has no effect on IL-10 during primary *Leishmania major* infection.**

Groups of WT and PTX3^-/-^ mice were infected with *L. major* and at the indicated times, (n = 5 per each time point), sacrificed and single cell suspension of the spleen and dLNs analyzed directly *ex vivo* by flow cytometry for the frequency of IL-10 producing CD4^+^ T cells (A, B & D). Splenocytes and dLN cells were also stimulated with SLA (50 μg/ml) for 72 h, and the levels of IL-10 (C & E) in the cell culture supernatant was measured by ELISA. Results are representative of three independent experiments with similar results. *, *p* < 0.05; **, *p* < 0.01.

**Supplementary Fig 4: rPTX3 inhibits excessive Th17 differentiation by purified CD4^+^ T cells from PTX3^-/-^mice.**

Highly purified CD4^+^T cells from WT and PTX3^-/-^ mice were labeled with CFSE dye and stimulated with soluble anti-CD3 and anti-CD28 antibodies under Th17-polarizing conditions in the presence or absence of different concentration of rPTX3. After 72 h, the frequency of IL-17A-secreting CD4^+^ cells was determined by flow cytometry. Representative dot plots (A) and bar graphs (B) showing the percentage of CFSE^-^ IL-17A^+^ CD4^+^ T cells. The levels of IL-17A in the culture supernatant fluids were assayed by ELISA (C). Results are representative of three independent experiments with similar results. *, *p* < 0.05; **, *p* < 0.01; ****, *p* < 0.0001.

**Supplementary Fig 5: PTX3 negatively regulates Th17-specific transcription factors.** Purified CD4^+^ T cells from WT and PTX3^-/-^ mice were stimulated with soluble anti-CD3 and anti-CD28 antibodies under Th17-polarizing conditions in presence or absence of rPTX3 (200/mL). After 72h, total RNA was isolated and mRNA levels of IL-17A (A) and RORγT (B), STAT3 (C), IRF4 (D), BATF (E) and AhR (F) were determined by RT-PCR. ***, *p* < 0.001; ****, *p* < 0.0001

**Supplementary Fig 6: rPTX3 treatment inhibits IL-17A production by spleen cells.** Groups of WT mice were with infected with *L. major* and at 24 hours post-infection, mice were divided into two groups (6 mice per group) and treated intralesionally with either PBS or rPTX3 (0.5 μg/mouse) 3 times weekly. At 4 weeks post-infection, mice were sacrificed and spleen cells were isolated and assessed directly *ex vivo* for the frequency of −17A (A) and IFN-γ (B) -producing CD4^+^ T cells by flow cytometry. Some spleen cells were stimulated with SLA (50 μg/ml) for 72 h, and the levels of IL-17A (C) and IFN-γ (D) in the cell culture supernatant fluids were determined by ELISA. Results are representative of two independent experiments with similar results. \**p* < 0.05, \*\**p* < 0.01.

**Supplementary Table 1: Clinical characteristics of Human CL patients**

